# Prediction Is Preserved but Long-Timescale Benefits Are Reduced in ADHD

**DOI:** 10.64898/2026.03.18.712582

**Authors:** Noam Tzionit, Danielle G. Filmon, Talia Maeir, Sage E. P. Boettcher, Anna C. Nobre, Nir Shalev, Ayelet N. Landau

## Abstract

Attention-deficit/hyperactivity disorder (ADHD) has been associated with atypical temporal processing across multiple cognitive domains. However, most evidence derives from simplified paradigms that isolate timing from spatial behaviour. Here, we examine how temporal prediction operates within a continuous, dynamic visual environment. Using the Dynamic Visual Search (DVS) task, we embedded spatiotemporal regularities into a sustained stream of visual events, allowing observers to implicitly learn and anticipate predictable targets. Continuous mouse tracking provided a fine-grained measure of action planning beyond discrete reaction time and accuracy metrics.

Young adults diagnosed with ADHD (N=40) were compared to matched neurotypical controls (N=38). Both groups benefited from target predictability and reduced distractor load, indicating intact early spatiotemporal learning in ADHD. Across the duration of the task, however, the groups diverged. Neurotypical participants showed progressive increases in behavioural benefits from prediction, accompanied by increasingly direct and efficient mouse trajectories. In contrast, individuals with ADHD reached a plateau in prediction benefits midway through the experiment. Their performance remained stable, with minimal evidence of resource depletion, but did not show further optimisation based on learned regularities.

These findings suggest that while prediction formation is preserved in ADHD, its progressive utilisation across longer timescales is attenuated. Rather than reflecting a primary deficit in learning or sustained attention, ADHD may involve altered long-timescale integration or weighting of predictive information in dynamic environments.

## Introduction

Attention-Deficit/Hyperactivity Disorder (ADHD) is a prevalent, multifaceted syndrome characterised by symptoms of inattention, hyperactivity, and impulsivity ^1^. It is estimated that about 5.3% to 7.2% of the general population is affected ^2,3^, decreasing to 2.5% in adulthood ^4^. Although ADHD is defined based on behavioural symptoms, it is often accompanied by atypical cognitive performance across multiple cognitive domains, including working memory, inhibitory control, attention, and more ^5–14^.

Theoretical accounts highlight timing-related deficits as a core component of atypical cognition and performance in ADHD. For example, Barkley (1997) considered difficulties in time representation as associated with behavioural inhibition deficits, limiting the ability of people diagnosed with ADHD to plan future goals and exercise self-control ^15^. Another theory discusses delay aversion in ADHD, a characteristic disproportionate discounting of time-delayed rewards ^16^. Similarly, it was suggested that limited temporal foresight, i.e., representing the consequences of one’s decisions and actions in the future, is affected in ADHD ^17,18^. In a comprehensive review paper, Noreika, Falter, & Rubia (2013) present multiple lines of evidence linking such temporal abnormalities directly to inattention and impulsivity symptoms.

Atypical time processing in ADHD is also reflected in basic cognitive functions. Individuals diagnosed with ADHD often struggle with tasks relying on implicit time representation across multiple scales. Short-term preparatory processes such as alertness ^8,19,20^, motor preparation ^21–23^, reward processing ^24–28^, and temporal expectations ^29^ are considered less efficient. For example, Dankner et al. 2017 have shown a reduced ability to inhibit saccadic eye movements before the appearance of a temporally predictable target in young adults diagnosed with ADHD ^29^. Such findings are taken as evidence of impaired ability to use implicit temporal structures to guide attention. In other work, ADHD has been associated with diminished performance gains from alerting signals preceding a target (^8,19,30^).

In addition to short-term temporal processes such as alertness and expectations, cognitive performance is also influenced by slower time-dependent functions, like sustained attention, vigilance, and arousal (e.g., ^31–33^). A substantial body of evidence links ADHD to difficulties in maintaining an adequate level of performance over extended periods. Theoretical models of ADHD, such as the cognitive-energetic model, specifically emphasise atypicalities in activation and effort – concepts closely related to arousal ^34–36^. Sustained attention capacity, defined as the ability to remain engaged with repetitive tasks over time, is frequently reported to be reduced in individuals with ADHD ^9,10,15,37^.

Most empirical evidence regarding temporal deficits in ADHD is confined to relatively simple task designs that maintain spatial elements constant. This methodology enables researchers to isolate the control of temporal behaviour but overlooks the reality that, in everyday scenarios, anticipating the timing of events is intrinsically linked to spatial behaviour (e.g., the school bus example). While research suggests that implicit spatial learning remains intact in ADHD ^38^, it is unknown if this extends to complex spatiotemporal regularities. Since temporal expectations play a crucial role in spatial guidance, core timing deficits in ADHD could disrupt performance in dynamic environments. Consequently, a crucial step forward involves investigating how timing deficits interact with the control of spatial attention

To address this, we utilised the Dynamic Visual Search (DVS; ^39,40^). This task assesses how well individuals learn and utilise spatiotemporal regularities embedded within a dynamic display where targets and distractors fade in and out. These events unfold in a continuous, extended stream rather than discrete trials, simulating the flux of visual information characteristic of real-world environments. Half of the targets are predictable in their onset time and location, while the others appear randomly. Previous studies employing the DVS have demonstrated that healthy observers, ranging from young children to older adults, implicitly learn to anticipate these targets, resulting in significant behavioural advantages ^40–43^.

In the current study, we applied this paradigm to compare young adults diagnosed with ADHD to age-matched, typically developing controls. We investigated the interplay of spatial selective attention and implicit timing mechanisms by manipulating goal-directed attention (via varying distractor set sizes) and spatiotemporal prediction (by embedding predictable targets). Furthermore, we traced the trajectory of performance across the duration of the task to evaluate learning and sustained attention over time. Crucially, we introduced continuous mouse-tracking analysis as a novel outcome measure to capture the full dynamics of action planning. While discrete metrics like reaction time provide a snapshot of performance, continuous trajectories offer a window into the real-time unfolding of motor control. This allows us to distinguish specific patterns of behaviour, revealing not just whether, but how attentional predictions dynamically reshape the guidance of action in ADHD compared to controls

## Results

### Behavioural Data

#### Reaction times

Analysis of the reaction times for finding targets in the dynamic visual-search (DVS) task revealed main effects of Group (β = .230, SE = .059, t = 3.888, p < .001, 95% CI [.114,.346]), Predictability (β = .2, SE = .016, t = 12.440, p < .001, 95% CI [.168,.231]), Distraction Load (β = .022, SE = .006, t = 3.857, p < .001, 95% CI [.011,.033]), and Block (β = -.081, SE = .006, t = -14.204, p < .001, 95% CI [-.092,-.069]). The control variable, Inter-Target Distance factor, was also significant (ITD; β = .025, SE < .001, t = 53.003, p < .001, 95% CI [.024,.026]). This significant main effect was a result of faster responses to targets that were closer to the location of the previous ones. As illustrated in Figures 2a and 2b, the four main effects were driven by faster reaction times in the group of neurotypicals compared to ADHD, when targets appeared predictably, when there were fewer distractors, and with progressing blocks.

Significant two-way interactions occurred between Group and Block (β = .029, SE = .008, t = 3.582, p < .001, 95% CI [.013,.044]) and between Predictability and Block (β = .033, SE = .008, t = 4.019, p < .001, 95% CI [.017,.048]). There was also a three-way interaction of Group X Predictability X Block. (β = -.023, SE = .011, t = -1.978, p = .048, 95% CI [-.045, -.0002]). The rest of the interactions were not significant (all p’s > .403).

To guide the interpretation of the interactions, we plotted the difference in reaction times for predictable vs. random targets for each experimental group as a function of experimental block (Figure 2c). The data indicated that the that the benefits in RT continued increased more steeply across blocks for NT compared to ADHD groups. The difference in the pattern of expectation-based benefits was validated using a post-hoc analysis to evaluate the linear trend, comparing the slope of the difference score to zero. In the control group, RT benefits increased across blocks with a negative slope (β = -.013, SE = .003, p < .001, 95% CI [-.02, -.007]). In the ADHD group, the slope did not differ significantly from zero (β = -.003, SE = .003, p = .323, 95% CI [-.009, .003]), meaning the difference did not change over time. The two slopes were also significantly contrasted between groups (slope difference = –0.01, SE = .005, t(310) = -2.163, p = .031), confirming that the learning progression differed between neurotypicals and the ADHD group.

#### Accuracy

Analysis of the accuracy for finding targets in the DVS task revealed main effects of Group (β = -.425, SE = .105, *z* = -4.03, *p* < .001, 95% CI [-.631,-.218]), Predictability (β = -.399, SE = .04, *z* = -9.989, *p* < .001, 95% CI [-.478,-.321]), and Block (β = .161, SE = .032, *t* = 5.073, *p* < .001, 95% CI [.099,.223]). There was no main effect of Distraction Load (p = .361) or any significant interaction (all p’s > .119). The main effects were driven by higher accuracy in neurotypicals compared to ADHD, when targets appeared predictably and with progressing blocks. The control factor of inter-target distances (ITD) exerted a main effect (β = -.033, SE = .001, t = -22.210, p < .001, 95% CI [-.035, -.030]) with targets closer in space to a previous target enjoying better accuracy.

#### Mouse Movements

We analysed mouse movements using the statistical contrasts described in the reaction-time and accuracy sections above. The data were compared using two complementary approaches. First, we calculated the average distance between the mouse pointer and the target at the time of the target onset by averaging the raw data. Importantly, using the gradual stimulus presentation, targets are, in fact, invisible at time of onset and only gain visibility as they increase intensity (the ramp, see black opacity line beneath mouse trajectory time series, Figures 3a, 4a, 5a, 6). In a second analysis, we aligned the data to response times, temporally locking the mouse trajectory at the moment of target selection (mouse click). The two approaches are complementary and emphasize different parts of the sensory-motor processing in the task. The target-locked analysis documents the initiation of mouse approach as sensory evidence accumulates. Since targets are invisible at onset time, this portion could express changes in performance due to the ability to spatially and temporally anticipate predictable targets, compared to random targets. The response-locked analysis allows a clean view of motor trajectories towards response as well as differences in response spatial precision (i.e., where or how near to the targets does the participant click). The two approaches are commonly used when investing trial-based time series averages (e.g., stimulus and response-locked ERPs).

Generally, target-locked mouse trajectories demonstrate a stable baseline, average distance from the target. Following almost 2 seconds, a marked reduction in distance to target reflects the average approach movement towards the target, reaching a minimum approximately 2 seconds after target onset, followed by an increase in the distance from the target, most likely reflecting the switching to the next target in the continuous, multi-target display. In describing our results, we refer to the first phase, leading to the time series trough as the approach phase and the subsequent phase, the switch phase. When comparing target-onset locked mouse trajectories between groups, we found a difference in the movement trajectory which was significant during the switch phase of the trajectory (i.e., following the distance trough) ([2233.33, 3650] ms, sum-t = 289.496, p = .004). This pattern indicates that, on average, participants in the control group moved away from the selected target (and on to the next) earlier compared to the ADHD group and aligns well with our behavioural data, indicating significant differences in mean RTs (Figure 3a).

Response-locked mouse trajectories demonstrate a pronounced trough around locking time (time zero). This is expected due to the inclusion of data from successfully detected targets only. Targets were labelled as detected if the participant clicked within a square window of ±0.90° visual angle (≈ 1.79° × 1.79°) centred on the target location and within the temporal window of target presentation (lasting 3.55 s). We found a significant difference between groups starting about one second before response time ([-1000, -466.67] ms, sum-t = 90.293, p = .025). The pattern indicated that the NT group took a shorter time to move the mouse and to select the targets they identified (Figure 3b). Phrased differently, the NT group traversed a larger distance within that fixed time window suggesting more efficient motor target selection routines.

In addition, there was a significant difference in mouse movements when comparing the raw distance between the mouse and the targets based on their predictability status. In the target-onset locked analysis, the mouse was significantly closer to the target for predictable target compared to random targets, during the approach phase of the movement (i.e., before and during the minima; [1033.33, 2383.33] ms, sum-t = -670.752, p < .001). Following the approach phase, a significant difference between predictability levels indicated a larger distance from predicted targets ([2533.33, 4000] ms, sum-t = 833.914, p < .001). This pattern suggests an overall more efficient approach and switch for predicted targets. This is displayed in Figure 4a. The interaction of Group and Predictability was not significant (p’s >= 0.181). When locking the signal to responses, we see that during the approach phase, there was significant difference between predictable and random targets ([-700, 333.33] ms, sum-t = -475.985, p < .001,). Also, there was a significant difference after response, showing a larger distance from predicted targets ([483.33, 1000] ms, sum-t = 158.909, p < .001). This is displayed in Figure 4b.

Although the response-locked analysis revealed two significant temporal clusters, the first clusters’ visual separation in the averaged trajectories was relatively subtle. To improve interpretability and provide a clearer sense of the magnitude and direction of the effect, we visualize the cluster effect using a box plot, including individual participants data.

A significant difference was also found when comparing mouse movements based on the number of distractors (high vs. low distraction load). Participants’ distance to target when there was low distraction load was smaller compared to when there was high distraction load ([1133.33, 2550] ms, sum-t = -309.578, p < .001, Figure 5a). The interaction effect of Group X Distraction Load was not significant (p’s >= 0.022).

When mouse trajectories are locked to response times, we observed a significant difference after response between distraction loads, where mouse distance to target was smaller when the number of distractors was smaller ([333.33, 1000] ms, sum-t = -126.534, p = .002, Figure 5b).

As the behavioural report (reaction times) above demonstrates, we also had significant two- and three-way interactions (Group by Block and Group by Block by Predictability, respectively). To examine these interactions in mouse trajectories we analysed the difference in mouse trajectories for predictable vs. random targets as a function of experimental block for each group separately. We subtracted the time course of mouse distance to random target from the time course of mouse distance to predicted target. For each group, this subtraction is performed per block separately in order to track those differences over the course of the experiment.

While analysing the signal relative to target onset, we observed an increase in the difference between predicted and random over time only for the neurotypical group, such that the benefit of predictable targets, over random ones, continued to increase over the experimental blocks. This benefit is depicted by an earlier start, closer minima to the target, and in the shorter overall time to perform the task. In contrast, the ADHD group did not show an increased benefit for predictable targets, over random targets, over time (Control: [816.67, 1766.67] ms, sum-t = 267.753, p = .001; ADHD: p’s >= .366). These results are presented in Figures 6a and 6b. When locking for response, we did not find any significant differences between the time-courses (p’s >= .083).

## Discussion

Participants responded faster when searching for targets among fewer distractors, when targets were predictable, and as the task progressed across blocks. Comparisons between the control group and participants with ADHD revealed that controls were overall faster and demonstrated an increase in prediction benefits over time. When comparing accuracy, all participants were more likely to find targets that appeared predictably and as blocks progressed. However, on average, the control group was more accurate than the ADHD group.

The control and ADHD groups differed in their ability to build stronger prediction benefits over time. While controls gradually increased their reliance on predictions as the task progressed, prediction benefits remained constant in the ADHD group.

Mouse□tracking data provide deeper insights into the dynamics of behaviour. Analysis of the raw (onset□locked) signals revealed that the primary group difference emerged in the post□response window (i.e., after the mean RT), a pattern consistent with differences in reaction times (earlier responses permit earlier disengagement from the selected target). However, when accounting for these differences by locking the signal to the response, it became clear that participants with ADHD took longer to move the mouse pointer to the target than controls. This suggests that slower reaction times largely reflect slower movements towards targets (i.e., less efficient motor behaviour).

Differences in efficiency were also evident when comparing movements towards random versus predictable targets. Before the response, trajectories towards predictable targets showed earlier reductions in mouseto□target distance (participants closed in on the target sooner). In contrast, for the two distraction□load levels, we did not observe such a pre□response pattern. Instead, in the response□locked analysis, the only significant difference between the load conditions occurred after target selection, reflecting post□selection processes. Overall, these results indicate that predictability supports more efficient control over the approach to the target.

Group differences in mouse-trajectories data were in line with the three□way Group × Block × Predictability interaction we observed when analysing reaction times. It was driven by changes in the magnitude of the difference between responses to predictable and random targets across experimental blocks. While the control group showed a gradual increase in the benefit from predictions over time, the ADHD group maintained a stable benefit throughout the task. Statistical analyses confirmed that this effect was restricted to an approximately 1□s window: in controls, the difference signal between trajectories to predictable and random targets increased significantly between ∼1 and 2 s after the onset of target fade □ in as blocks progressed, whereas the ADHD group showed no significant change at any time point.

Our findings replicate and extend previous work on dynamic visual search. As in prior studies, predictions conferred significant benefits ^40–42,44^, and adding distractors impaired performance^55^. In the current study, we focused on ADHD, a group that often shows considerable differences compared to age-matched controls in experimental tasks involving temporal attention ^11,24,29,56^. While replicating previous findings of overall slower responses in ADHD (e.g., ^47,48^), we identified a more nuanced group difference. The ability of individuals with ADHD to use spatiotemporal regularities is constrained: they utilise predictions early in the task but do not increase their reliance on learning over time in a way that appears adaptive. This lack of progression may limit their ability to gradually free up important attentional resources. This pattern was evident in both mean reaction times and response dynamics.

The analysis of mouse movements provides meaningful insights into the performance benefits conferred by attention. Hypothetically, faster responses can arise from various reasons. For instance, participants might use predictions to initiate mouse movements earlier towards a target without altering their movement speed. Conversely, participants might use their knowledge of spatiotemporal regularities to adopt a different strategy—spending more time exploring the rest of the display and compensating by moving faster towards the target. Our findings showed that faster movements were characteristic of selecting a predictable target, as well as a key source of the main effect of the Group factor. In contrast, the main effect of task difficulty was not associated with faster mouse movements. These differences in motor trajectories may reflect different sources of attentional guidance, with experience-driven components supporting more proactive behaviours.

In line with previous studies, we observed that ADHD did not show a specific deficit in top-down control, which was manipulated by increasing the number of distractors. Instead, we replicate previous findings indicating differences in mean performance indices (e.g.,^47,48^), potentially due to delayed early perceptual processing and motor selection (e.g.,^50^). The dynamic markers provide an interesting insight: on average, the ADHD group started moving later towards targets compared to the control group, and their movement took longer.

Our findings on the limited utilisation of prediction add an important piece to the puzzle of attention control in ADHD. First, we observe that students diagnosed with ADHD demonstrate impressive learning abilities, as reflected in their prediction utilisation across behavioural markers. Previous studies have reached similar conclusions, suggesting that the core mechanisms of visual statistical learning remain intact in ADHD (e.g., ^57–59^). A unique aspect of our study is the inclusion of time-related learning, which may pose specific challenges for individuals with ADHD ^29^. Our experimental design juxtaposes two opposing factors influencing behaviour in ADHD: intact learning mechanisms alongside temporal deficits. While we found that implicit learning of spatiotemporal regularities was present in the ADHD group, we did not observe the same increase in benefits over time as in the control group.

Sustained attention deficits, which are commonly observed in ADHD ^15^, cannot fully explain the significant interaction pattern. Although poor sustained attention affects performance in repetitive tasks ^33,60^, our behavioural data revealed improvement trends across successive blocks in both groups. This suggests that the differences are not purely “energetic” in nature. Such a pattern, on its own, may not be surprising as our task was very different to traditional designs of sustained attention, which are inherently less engaging ^60–63^.

Although previous studies examining temporal expectations in ADHD argue for a deficit in short-term preparatory processes (^8,19,20,30,56^), our results highlight an interesting refinement. The ADHD group could anticipate predictable targets, as revealed in the substantial behavioural benefits irrespective of group. They also showed limited or no signs of global fatigue in their performance on the task (e.g., over RT and accuracy stable). However, utilization of the learned predictions was exhausted midway through the experiments’ duration. The impact of the passage of time, operationalised as block number, affected the groups differently. The behavioural pattern reveals a failure to maximise the predictive utility over time in the ADHD group. To the best of our knowledge, previous studies that investigated temporal deficits in ADHD did not incorporate how those may evolve over time, and instead focused on mean differences. It is possible that our task managed to capture a unique property of cognition in ADHD that is revealed in an extended context. Such a trend aligns with broader literature characterising ADHD as a syndrome of dysregulated thought and action that diminishes optimisation of learned regularities. ^64^

To conclude, we present novel data elucidating fundamental cognitive mechanisms and the distinctive characteristics of ADHD during complex, dynamic tasks. By analysing behavioural markers and continuous recordings of mouse movements, we demonstrate how observers search for fleeting targets, extract spatiotemporal regularities, and manage varying levels of distraction. Mouse tracking further revealed distinct performance factors that underpin these processes, including enhanced efficiency across the entire response trajectory. When comparing control participants to those with ADHD, we observed divergent trajectories in prediction-guided attention over time: controls exhibited typical, adaptive learning that facilitates proactive guidance of attention in both time and space, whereas individuals with ADHD showed a pronounced limitation in capitalising on task utility over extended periods. A possible interpretation for this pattern might point to motivational or reinforcement-related mechanisms that dynamically increase the weighting of predicted information. Altered valuation of predictions could contribute to the reduced long-timescale optimization observed in ADHD. Together, these findings represent a significant advance in our understanding of cognitive control in dynamic, real-world scenarios, where anticipatory signals and current goal sets jointly shape both attention and motor behaviour.

## Methods

This study was reviewed and approved by the Faculty of Social Sciences ethics committee at the Hebrew University of Jerusalem (approval number 2023-09212).

### Participants

The sample size was determined based on previous research^39,40,43^. A simulation-based power analysis using the mixedPower Package in R ^65^ indicated that a sample size of 25 participants would lead to a power above 80% for detecting an effect that is half as large as the one we observed in the data obtained by ^40^. However, due to the use of new indices (mouse tracking) and the inclusion of cognitively diverse individuals, we increased the sample to a minimum of 40 participants (representing a 60% increase).

The original sample consisted of 46 control participants without ADHD diagnosis and 49 ADHD-diagnosed participants who provided written consent for course credit or monetary compensation (50 NIS/∼13 USD). All participants had normal or corrected-to-normal vision and no history of psychiatric or neurological disorders (excluding ADHD for the ADHD group). ADHD participants indicated that they exclusively used methylphenidate-based medications and abstained from taking their ADHD medication on the day of the experiment. Exclusion criteria were based on participants’ failure to complete the study procedures, missing mouse data, or poor performance (exceeding 2 SD compared to the mean of each group).

In the control group, out of 46 participants (*M*_*age*_ = 24.3, *SD*_*age*_ = 5.39; 73.91% females), 38 were included in the analyses; 1 incomplete recording; 2 missing mouse data; 5 participants with a performance below 2 SD from the mean performance of all subjects were discarded. In the ADHD group, out of 49 ADHD participants (*M*_*age*_ = 24.56, *SD*_*age*_ = 2.59, 57.14% females), 40 were included in the analyses; 6 incomplete recordings; 3 participants with a performance below 2S D from the mean performance of all participants were discarded. A detailed description of group demographics after exclusion is provided in Table 1.

**Table 1:**
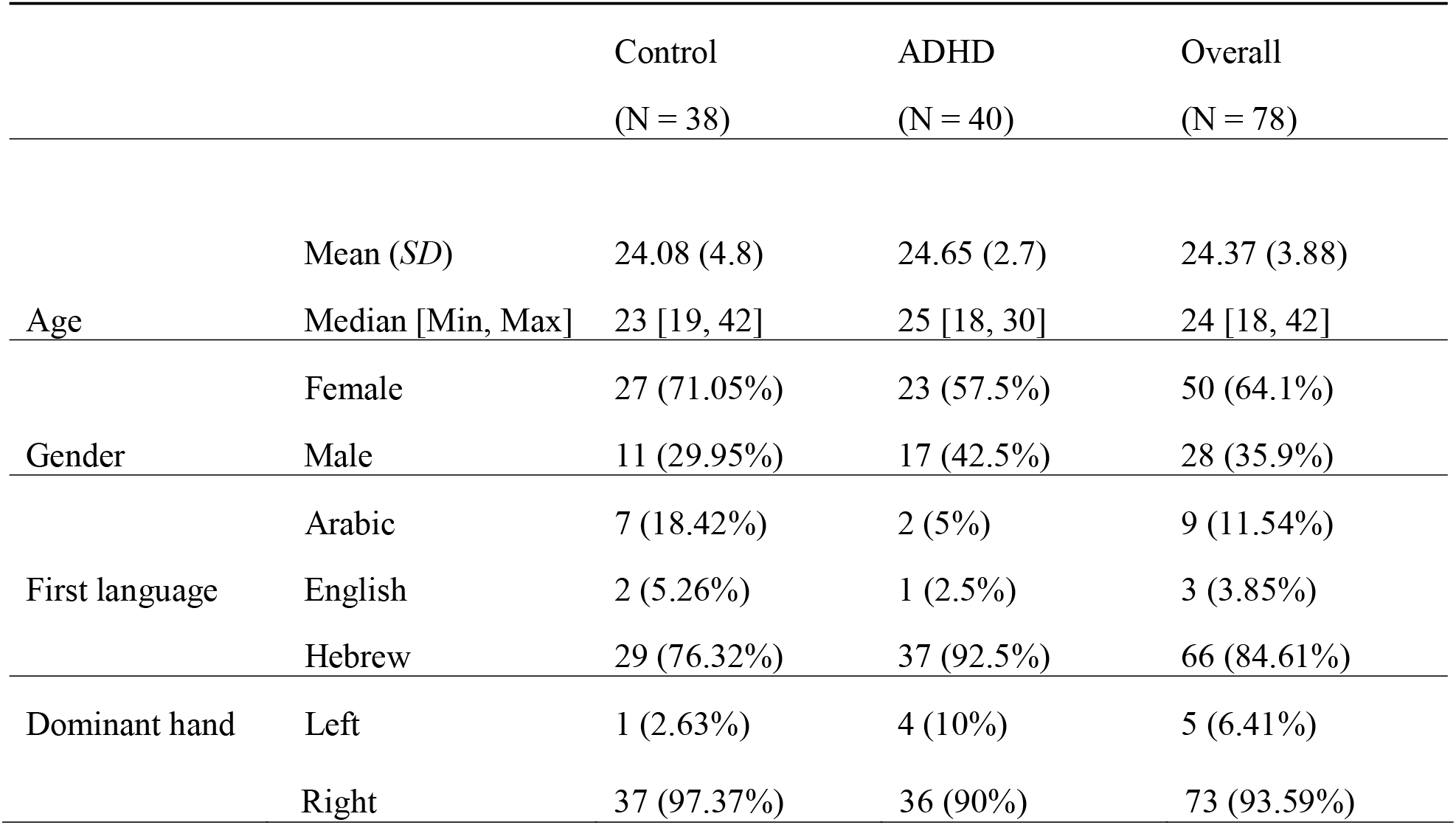
Demographic Characteristics of Study Participants by Groups. SD = standard deviation. There were no statistically significant differences between the groups.

#### ADHD Diagnosis

Participants in the ADHD group had undergone a formal diagnosis prior to and independent of the present study, conducted by a certified neurologist or psychiatrist following Israeli Ministry of Health guidelines. The comprehensive clinical assessment includes a detailed history, academic performance, medical conditions, family history, and previous psychological evaluations. Clinicians assess how ADHD symptoms manifest in different life contexts. This thorough assessment ensures a consistent pattern of attention challenges over time, aligning with DSM-5 criteria and involving objective tests and clinical interviews.

### Apparatus

Participants performed the DVS task^31,32,46^ (Figure 1). The search array consisted of oriented lines that ebbed and flowed into a dynamic display. The target was predefined as the vertical line, and distractors were slanted lines. Each trial contained eight targets: four predictable and four random.

**Figure 1.**
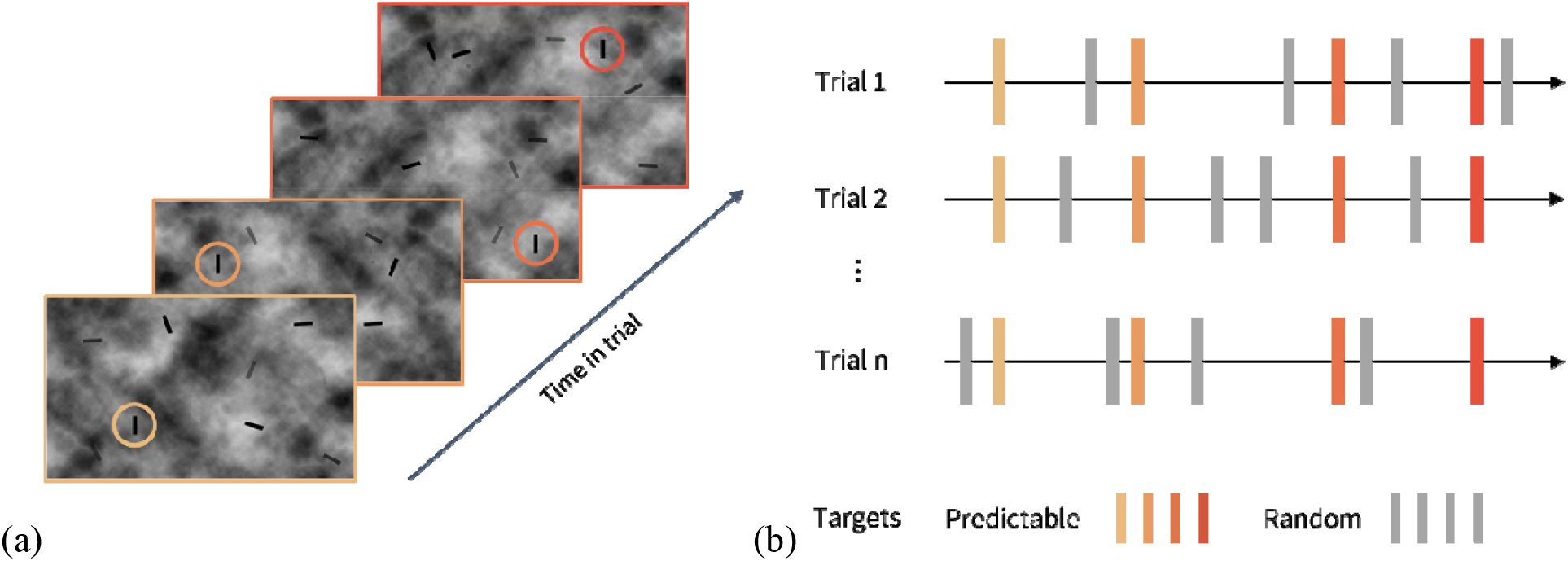
The Dynamic Visual Search Task (DVS). (a) An illustration of the experiment display and stimuli as a function of time in a single trial. Elements in the display appeared briefly (∼1 s fade in, ∼1.3 s persistence on the screen, and ∼1 s fade out). Targets were vertical lines, and distractors were slanted lines. Target onsets could be predictable (P) in time and spatial quadrant or random and thus unpredictable (U). Each trial lasted approximately 14 s and included four predictable and four random targets. Predictable targets were presented at a predictable moment within a given quadrant. Quadrants-time contingency (for predictable targets) is colour-coded in the illustration for clarity (from light to dark orange). (b) An illustration of three trials with target times indicated as vertical elements. Colours for predictable targets as in (a), random targets are coded grey.

**Figure 2.**
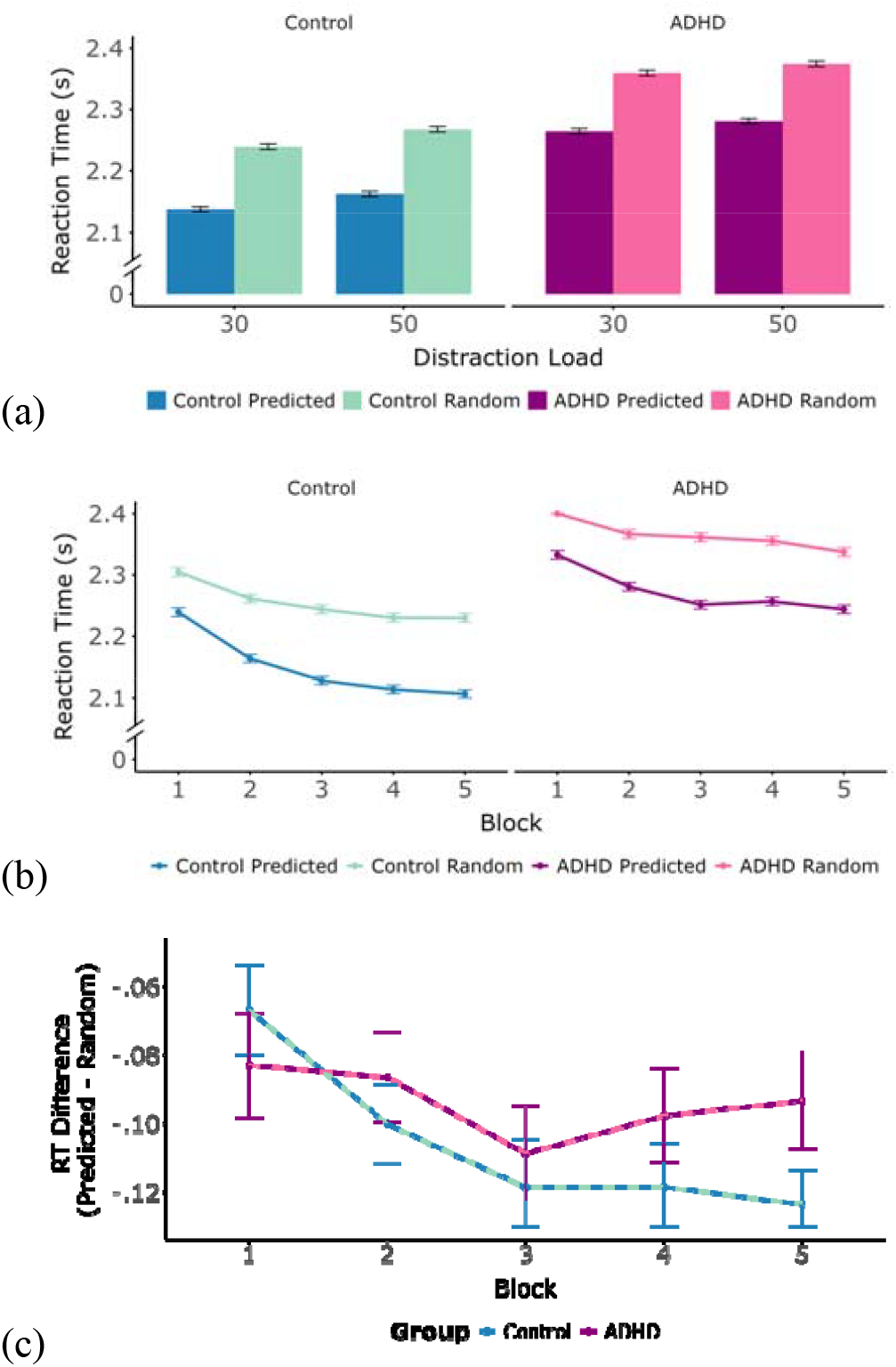
(a) Mean Reaction Times for each Group by Predictability and Distraction Load. The error bar represent ± 1 standard error (SE) of the mean reaction time. SE was calculated as the standard deviation of reaction times divided by the square root of the number of participants for each combination of group, predictability, and distraction load. (b) Mean Reaction Time for each Group by Predictability, acros Time in the Experiment. The error bars represent ± 1 standard error (SE) of the mean reaction time. SE was calculated as the standard deviation of reaction times divided by the square root of the number of participants for each combination of group, predictability, and block. (c) Differences in Mean RT for each Group across the Experiment Blocks. The error bars in the difference plot represent ±1 standard error (SE) of the reaction time difference between Predicted and Random targets. The SE was calculated based on the standard deviation of the reaction time differences (Predicted – Random) for each participant, divided by the square root of the number of participants within each group and block.

**Figure 3.**
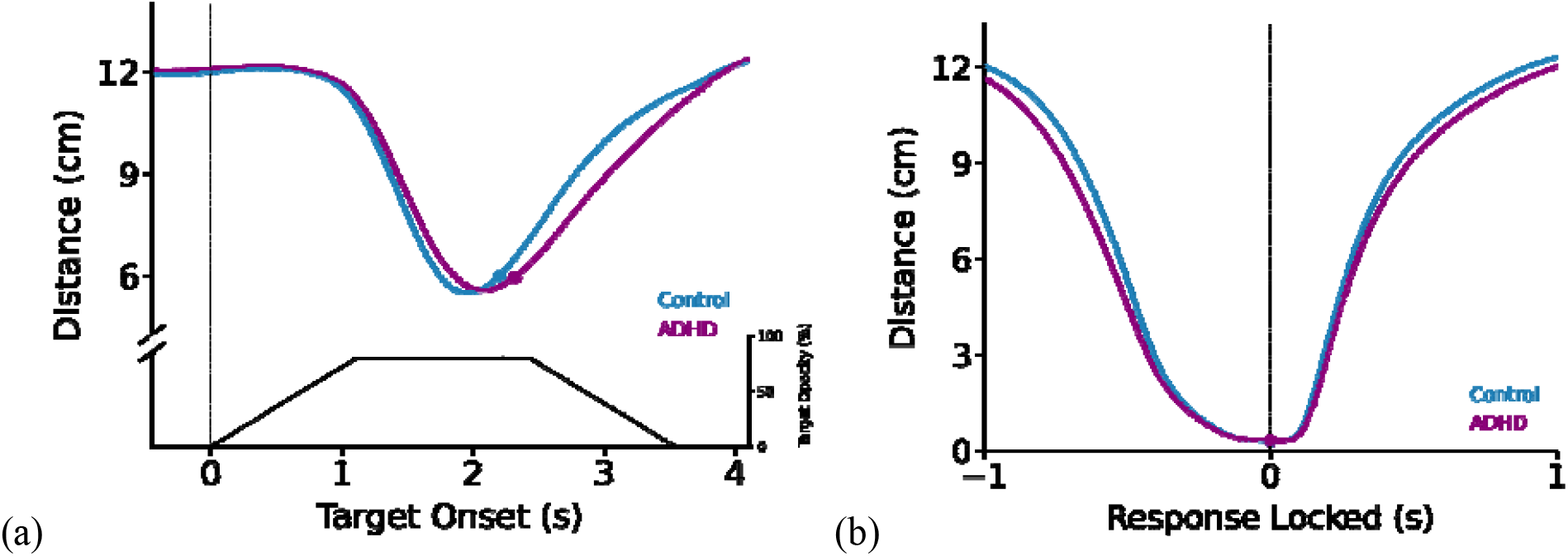
(a) Average Mouse Distance from Target by Group. This plot showcases the mouse distance in cm, locked to participants’ response times. The y-axis shows the mouse distance from the target in cm. The vertical dashed line indicates the target onset time. Shaded regions around the lines depict th standard error of each time point. Significant differences between the groups are highlighted in grey. Points on the time series indicate the average reaction time for the control group (blue) and the ADHD group (purple). Black line on the same plot indicates the opacity-time course including the ramp-on, stable and ramp-off phases of target appearance; the right-hand axis shows opacity in percent (0–100%), where 80% corresponds to the maximum target opacity. (b) Average Mouse Distance from Target by Group, Locked to Response. The x-axis represents the time window, spanning -+1 seconds relative to the response time. Shaded regions around the lines depict the standard error of each time point. The vertical dashed line indicates the time of the mouse click. Significant differences between the groups ar highlighted in grey. Points on the time series indicate the average reaction time for the control group (blue) and the ADHD group (purple).

**Figure 4.**
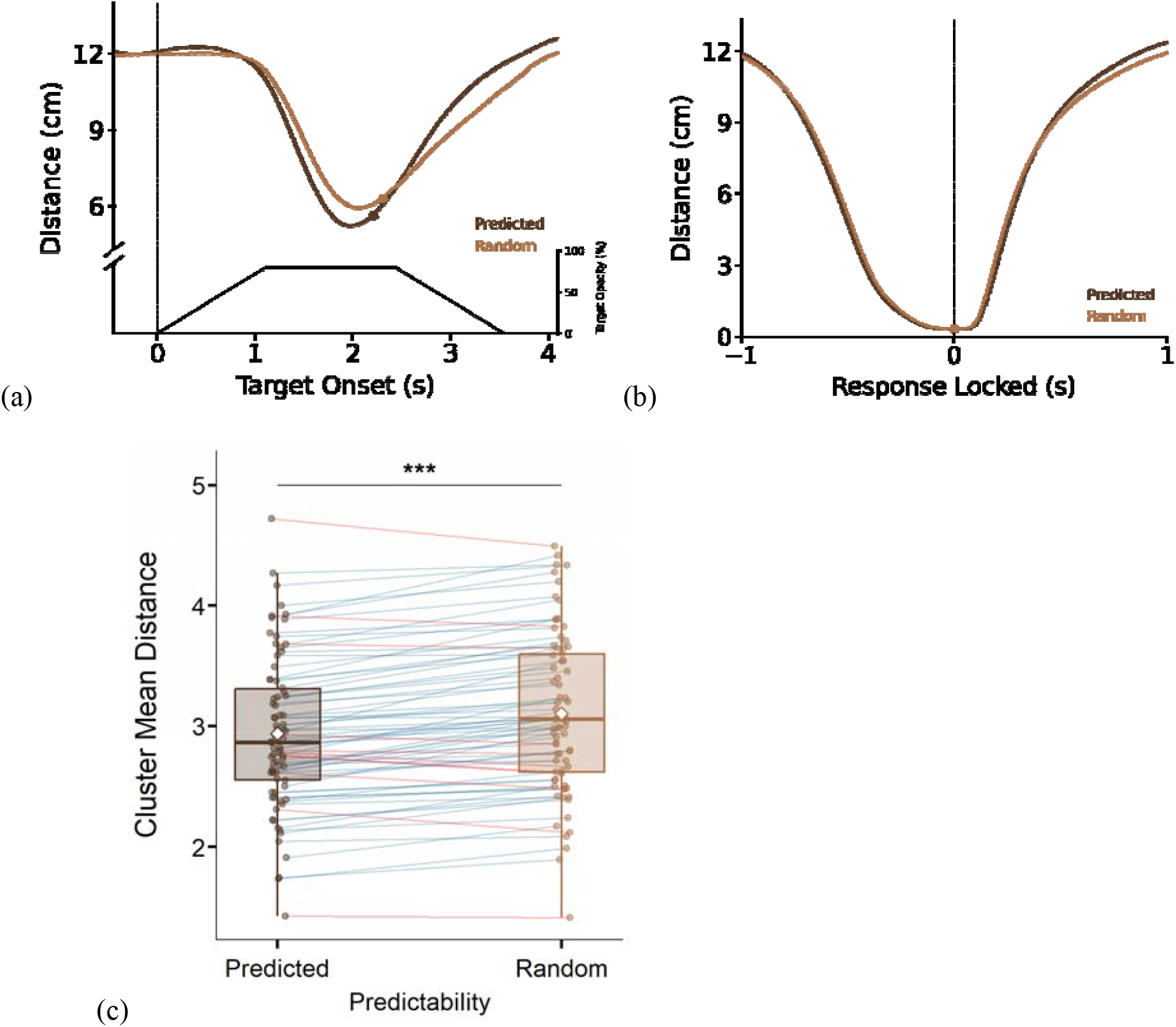
(a) Average Mouse Distance from Target by Predictability. This plot showcases the mouse distance in cm, locked to participants’ response times. The y-axis shows the mouse distance from the target in cm. The vertical dashed line indicates the target onset time. Shaded regions around the lines depict the standard error of each time point. Significant differences between predictability types are highlighted in grey. Points on the time series indicate the average reaction time for predicted (dark brown) and random (light brown) predictability. Black line on the same plot indicates the opacity-time course of target appearance; the right-hand axis shows opacity in percent, where 80% is the maximum opacity. (b) Average Mouse Distance from Target by Predictability, Locked to Response. The x-axis represents th time window, spanning -+1 seconds relative to the response time. Shaded regions around the lines depict the standard error of each time point. The vertical dashed line indicates the time of the mouse click. Significant differences between predictability types are highlighted in grey. Points on the time serie indicate the average reaction time for predicted (dark brown) and random (light brown) predictability. (c) Cluster Summary Comparison of Mouse Distance by Predictability. To complement the response-locked time-series analysis, this plot visualizes the magnitude and direction of the first significant cluster in the ‘Average Mouse Distance from Target by Predictability, Locked to Response’ plot (Figure 4b). The y-axis reflects the mean distance between the mouse and the target location, computed across the tim window of the first significant cluster. Each point represents a participant’s mean distance for the predicted or random targets. Lines connect paired observations from the same participant, illustrating within-subject differences. Blue lines indicate participants for whom predicted targets were closer (smaller distance) than random targets, whereas red lines indicate the opposite pattern, where predicted targets were farther than random targets. Boxplots summarise the distribution of cluster means for each condition (median, interquartile range, and whiskers), and diamond markers indicate condition averages.

**Figure 5.**
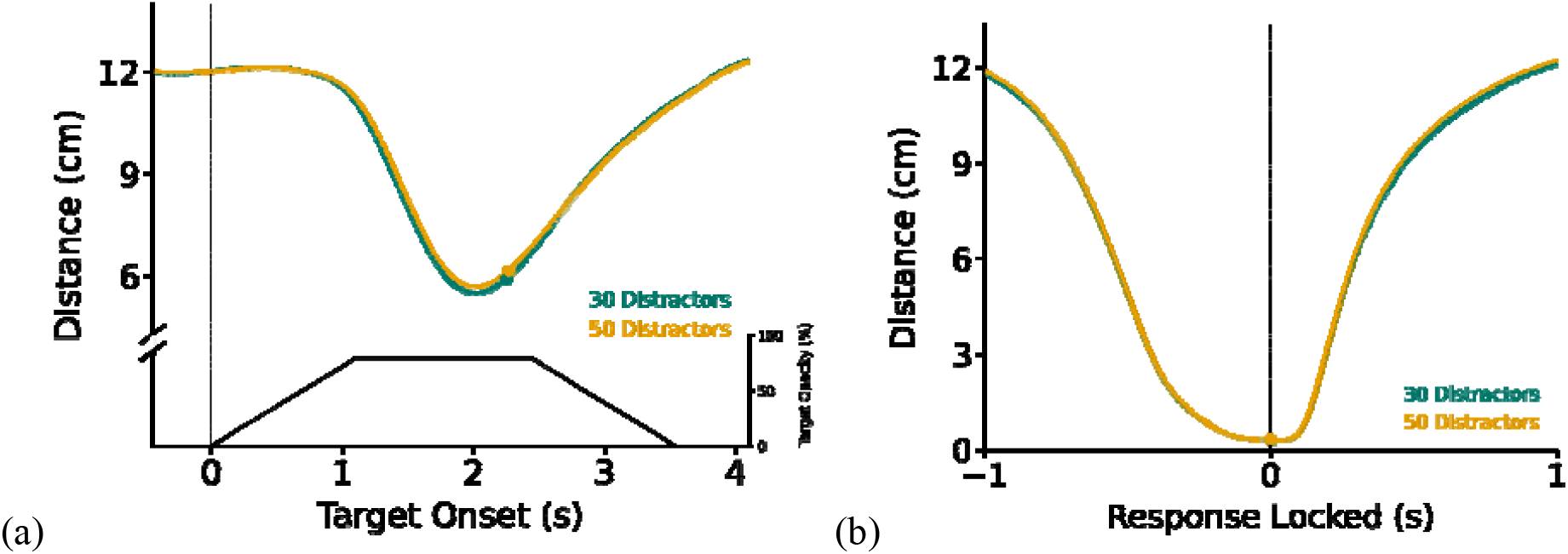
(a) Average Mouse Distance from Target by Distraction Load. This plot showcases the mous distance in cm, locked to participants’ response times. The y-axis shows the mouse distance from the target in cm. The vertical dashed line indicates the target onset time. Shaded regions around the lines depict the standard error of each time point. Significant differences between distraction loads are highlighted in grey. Points on the time series indicate the average reaction time for low (30; green) and high (50; yellow) loads. Black line on the same plot indicates the opacity-time course of target appearance; the right-hand axis shows opacity in percent, where 80% is the maximum opacity. (b) Average Mouse Distance from Target by Distraction Load, Locked to Response. The x-axis represents the time window, spanning -+1 seconds relative to the response time. Shaded regions around the lines depict the standard error of each time point. The vertical dashed line indicates the time of the mouse click. Significant differences between distraction loads are highlighted in grey. Points on the time series indicat the average reaction time for low (30; green) and high (50; yellow) loads.

**Figure 6.**
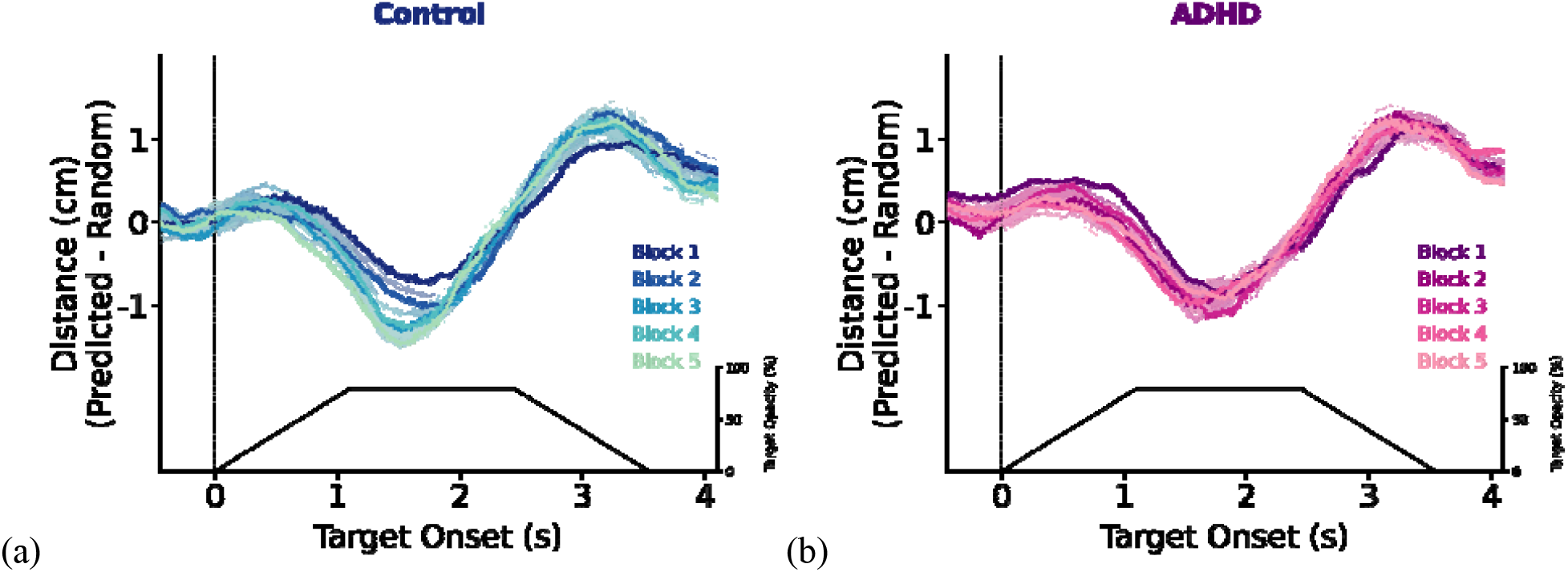
(a) Control Group Average Predictability Difference of Mouse Distance from Target by Block. This plot showcases the difference between mouse distance from predicted and random targets in cm, locked to target onset time. The y-axis shows the differences in mouse distance from the target in cm. Negative values indicate that the mouse in the predictable condition was closer to the target compared to the distance to random at that time point. The vertical dashed line indicates the target onset time. Shaded regions around the lines depict the standard error of the mean of differences per time point. Significant differences between blocks are highlighted in grey. Time-course colour (dark to light blue) indicate data from blocks 1-5, respectively. Black line on the same plot indicates the opacity-time course of target appearance; the right-hand axis shows opacity in percent, where 80% is the maximum opacity. (b) The same as (a) only for the ADHD Group Average Predictability Difference of Mouse Distance from Target by Block. This plot showcases the difference between mouse distance from predicted and random target in cm, locked to target onset time. The y-axis shows the mouse distance from the target in cm. Th vertical dashed line indicates the target onset time. Shaded regions around the lines depict the standard error of the mean of differences per time point. Time-course color (dark to light purple) indicate data from blocks 1-5, respectively.

Predictable target timings and locations were implemented following the dynamic search paradigm described in previous work (^40^, Experiment 2). Predictable targets appeared at the same time relative to the start of the trial and in the same quadrant on every trial. Four predictable target onset times were selected from six predefined temporal bins distributed across the trial. For a given participant, the four predictable target times were repeated throughout the experiment. Although predictable targets appeared at a predictable time within a specific quadrant of the display, their specific location within the quadrant was random.

In addition to the predictable targets, each trial included four random targets and either 30 or 50 distractors. We uniformly distributed the onset of these unpredictable items across the trial duration by pseudo-randomly choosing a start time for each item to begin to fade in. This distribution was constrained to present a quarter of the distracting items and targets within each successive 2.5-second window (thus creating 4 artificial and seamless time bins within the trial). Once the items were assigned a particular starting time, we chose four of those items to serve as the random targets.

This assignment was not contingent on the temporal distance between the unpredictable targets or the distance to the predictable targets. This means that the exact temporal relations between distractors and unpredictable and predictable targets were not predetermined or constrained. All stimuli had the same temporal profile, and a trial ended when the last stimulus had completely faded out of the display. The spatial locations of the distractors and unpredictable targets were determined randomly. For each item, we randomly chose a quadrant (1–4) and then, within that quadrant, randomly chose a spatial location, with the only constraint being that the items should not overlap or cross the quadrant border. Note that this means that the number of unpredictable targets in each quadrant was unpredictable on any given trial.

### Stimuli

The search display comprised a full field with four regions containing 1/f static noise patches generated uniquely for each trial and connected seamlessly to cover the entire screen (dimensions of 1920× 1080 pixels). Throughout each 14-s trial, either 30 or 50 distractor stimuli (bars tilted either left or right, with orientations ranging from 80°–100° and 260°–280°) and eight target stimuli (vertical bars, 0°) appeared and disappeared at different times.Targets and distractors were all dark grey (RGB values [40 40 40]) and gradually faded in and out over an epoch of 3.55 s. The stimulus ramped up over 1.11 seconds to reach maximal opacity (80%), remained at maximum visibility over 1.33 seconds and then faded out over 1.11 seconds. Each stimulus was 0.89° in length and 0.15° in width with no overlap with other stimuli. For analysis purposes, onset times were defined as the first moment when the target stimulus commenced fading in.

### General Procedure

After providing consent, participants completed a demographic questionnaire (reporting on items such as age and hand dominance) and received an explanation about the task and procedures. They were seated in a dimly lit room with their head resting on a chin rest and were asked to complete a short 4-trial training followed by the DVS task lasting approximately 45 minutes.

Participants were instructed to click using their mouse at the target location to indicate its detection. A total of 1,600 target events were presented across five blocks. Each block consisted of 40 trials (with 8 targets each). After each trial, observers received feedback in the form of a number between 0 and 8, indicating how many targets they found (e.g., “You found 5 targets out of 8”). The next trial was initiated by the participant (self-paced). In addition to including predictable and random targets, the current design also manipulated the number of distractors, introducing two levels of load. Th experimental procedure relied on the experimental code used by^40^, with the addition of the manipulation of the distractor load (The task is illustrated in Figures 1a & 1b).

### Behavioural Analysis

#### Accuracy and RT Analysis

Our behavioural analysis was designed to estimate group differences (ADHD versus neurotypical controls) in four within-subject factors that are relevant to our research objectives: target predictability (predictable versus random), distractor load (high versus low), and block number (the number of blocks between 1 and 5). In addition, Inter-Target Distance (ITD; a continuous variable) was considered as a control variable. This variable captures the distance in space between each pair of sequential targets and controls for this trivial source of variability in performance in the highly dynamic search displays.

We employed two types of models to analyse our data: Linear Mixed Models (LMM) for reaction times (RTs) and Generalized Linear Mixed Models (GLMM) for accuracy. The dependent variables: RTs and Accuracy Rate, were scaled and centred and treated as continuous. Both models initially included the maximal random-effects structure as recommended by ^66^. Random slopes that did not improve the model fit were removed based on likelihood ratio tests. The independent variables in both models were Predictability, Distraction Load, Time in the experiment, and Group.

The LMM included experimental conditions such as fixed and random effects for participants to account for individual differences. Initially, the random-effects structure included intercepts and by-participant slopes for the independent variables. The optimal model retained random intercepts for participants and slopes for Predictability. Additionally, we included a control variable reflecting the distance of each target from the previous target. For the first target in a trial, we calculated the distance between the position of the first target and the first recorded mouse position in each trial.

Accuracy was analysed using a GLMM, treating it as a binary outcome. The fixed effects included experimental conditions, and random effects were specified for participants. The maximal initial random-effects structure included intercepts and by-participant slopes for Predictability, Distraction Load, and Block in the experiment. The final model included intercepts for participants and slopes for Predictability, Distraction Load, and Block. We also included a control variable of the distance between each target and the previous one.

Significant interactions were explored using the emmeans package ^48^ with Tukey post hoc correction to adjust for multiple comparisons.

#### Mouse-tracking Analysis

Reaction times and accuracy measures provide discrete measures of target performance. To further capitalize on the ongoing, dynamic nature of the utilized task (DVS) we included an of mouse movement trajectories. This measure provides a continuous measure of the selection process that leads to target detection. Our comparisons rely on the same variables as with reaction times and accuracy: the influence of Group, Target Predictability, Distraction Load, and Block Number on mouse movements toward targets within a designated time window.

We sampled the position of the mouse pointer throughout each trial at a refresh rate of 60 Hz. This resulted in a continuous vector that represented the evolving motor behaviour of participants as they engaged in the task. Recording was done using a MATLAB^67^.

Raw mouse data were generated through the experimental MATLAB code. The mouse data matrix comprised X and Y coordinates of mouse position, click events, and indications of whether a target was detected (i.e., a hit). After combining the mouse data with the rest of the logged experimental data, the motor trajectories were processed by calculating the Euclidean distances between the raw mouse positions and the target location within specific time windows. This results in a time series delineating the evolution of mouse distance. We used a window spanning 500 ms before the target onset to 4000 ms after the target onset. To estimate the mouse trajectories uninfluenced by the substantial variance in reaction times, we locked the mouse trajectories to the moment of the response click on the target, using a window spanning from 1000 ms before the mouse click to 1000 ms after the click.

We first examined the main effects of each independent variable through four primary comparisons, which included Group (ADHD vs Control participants); Predictability (Predicted vs Random targets); Distraction Load (Low vs. High load); Block (blocks 1–5). To assess whether group differences modulated task performance, three interaction effects were examined, Group and Predictability, evaluating whether target predictability affected ADHD and Control participants differently; Group and Distraction Load, Testing whether the impact of distraction varied across groups; and Group and Block, investigating learning and adaptation effects in each group over time. Post-hoc analyses were conducted to explore and visualise interaction effects.

To test for statistically significant differences in the comparisons mentioned above and account for the multiple comparisons, a non-parametric cluster-based approach was used and implemented in the Fieldtrip toolbox^68^. Clusters were formed using neighbouring points, based on the Monte-Carlo method. Significance probabilities were calculated based on permuting the data 10,000 times between the experimental conditions. Following our hypothesis, the critical alpha level was fixed at 0.05 (two-sided). For significant clusters, we report the sum-t statistics (the sum of all t-values in the cluster) and p-values.

For the post-hoc comparisons, we applied the same cluster-based permutation test but introduced further corrections for multiple comparisons using the Bonferroni method. This conservative approach adjusts the alpha level by dividing it by the number of possible combinations between the involved variables, ensuring a stringent control of false positives. All cluster analyses were based on participant averages. Note that the number of targets between each experimental condition differs because we only included mouse movement for successfully detected targets.

## Notes

### Competing Interest Statement

The authors have declared no competing interest.

### Funding

- Israel Science Foundation: 1073/24, 1899/21, 958/16
- Academy of Medical Sciences: DTTFR13\1107
- European Research Council: 852387

